# KMT2C and KMT2D amplify GRHL2-driven enhancer activation

**DOI:** 10.64898/2026.03.10.710694

**Authors:** Ryan M. Boileau, Kevin X. Chen, Robert Blelloch

## Abstract

The activation of cis-regulatory enhancers is essential for cell fate specification by driving cell type-specific gene expression. Differentiation models are widely used to study enhancer biology but the asynchronous and interdependent nature of gene regulatory changes during cell state transitions can complicate mechanistic studies. To overcome these limitations, here we develop a tamoxifen-gated system for acute enhancer activation in embryonic stem cells (ESCs) based on GRHL2, a pioneer transcription factor which naturally becomes expressed as naive ESCs differentiate into the formative ESC state. Using this system, we investigate the functional relationship between GRHL2 and the histone mono-methyltransferases KMT2C and KMT2D (KMT2C/D). GRHL2 readily binds its target sites independent of KMT2C/D. However, in the absence of KMT2C/D, there are dramatic reductions in H3K4me1/2, P300 recruitment, and H3K27ac deposition at these sites as well as diminished transcriptional activation. Still, strikingly, a basal level of active enhancer mark acquisition and transcriptional activation occurs. Consistent with these findings, during the naive to formative ESC differentiation, GRHL2 enhancer remodeling and target expression is also strongly but incompletely dependent on KMT2C/D. Together, these results define a functional co-activator relationship in which KMT2C/D act as important amplifiers of GRHL2-driven enhancer activation in ESCs and establish a rapid inducible system for dissecting the kinetics and enzymatic dependencies of pioneer transcription factor mediated enhancer remodeling.

## Introduction

The acquisition and maintenance of new cell fates during development relies on the precise spatiotemporal regulation of gene expression. Central to this regulation is the cell-type specific expression of transcription factors (TFs), which bind to specific DNA motifs and function largely through the chromatin regulating complexes that they recruit. In addition to gene promoters, a major class of TF binding elements that influence transcription are distal cis-regulatory elements known as enhancers. The interplay between enhancers, TFs, and chromatin regulators is critical for converting an inactive enhancer to an active, transcriptionally promoting enhancer state. Disrupting enhancer state dynamics through loss- or gain-of-function mutations in any one of these classes of factors can have severe developmental and disease-related consequences^1,2^.

While only a few TFs may bind a particular enhancer, several key chromatin regulators are broadly associated with enhancer function. This includes the histone methyltransferases KMT2C/D (COMPASS-like complex) and histone acetyltransferases CBP/P300 which deposit H3K4me1/2 and H3K27ac respectively^3–6^. These histone modifications are considered to be active marks of enhancers and their deposition is predictive of enhancer activation^7–9^. A molecular epistasis for the early stages of enhancer activation from an inactive state has been proposed. Enhancer-bound TFs recruit KMT2C/D; KMT2C/D deposit H3K4me1/2 and recruit CBP/P300^7–9^; CBP/P300 deposit H3K27ac, recruit additional factors, and promote transcription^10,11^. Recently, multiple modes of enhancer activation have been demonstrated^12^, identifying both KMT2C/D -independent and -dependent mechanisms. One likely interpretation is that only certain TFs are dependent on KMT2C/D function, extending previous observations on TF specificity with other chromatin regulators^13^. Thus, the exceptions and rules to the models of enhancer activation must take into account the context of specific transcription factors.

A common system for studying TFs and enhancers is the naive to formative embryonic stem cell (ESC) transition. This *in vitro* differentiation highly recapitulates gene regulatory events that occur during mouse early embryogenesis in the epiblast between E4.5 and E5.5 including enhancer activation^14–17^. During the naive to formative transition the master epithelial TF Grainyhead-like 2 (GRHL2) becomes expressed and activates a few hundred enhancers to drive formative gene transcription and a mesenchymal-to-epithelial transition^18–21^. When prematurely expressed in naive ESCs, GRHL2 is also sufficient to bind and activate formative targets^21^ that are occluded in the naïve state, consistent with its role as a pioneering factor in facilitating chromatin remodeling to activate transcription^22^. Expectedly, loss of function of GRHL2 during development results in several defects including in neural tube closure and lethal craniofacial malformations^23–26^. Moreover, GRHL2 also functions as a suppressor of metastasis in a variety of epithelial derived cancers including gastric, lung, and breast cancers^27–29^. Therefore, identifying mechanisms of GRHL2 function is anticipated to provide both general insight into pioneer transcription factor function and the particular biology of this important developmental- and disease-related gene.

Herein, we engineered a conditional GRHL2 system for the study of acute enhancer activation in the context of naive ESCs. In this study, we interrogate the role of KMT2C/D for GRHL2 function. We show that synthetic translocation of GRHL2 to chromatin mediates H3K4me1 and H3K27ac deposition through recruitment of KMT2C/D and that KMT2C/D are required for GRHL2-driven transcriptional activation. Consistent with these findings, we demonstrate using formative ESC differentiation that KMT2C/D are required for expression of GRHL2 targets and facilitating the remodeling of GRHL2 enhancers with active enhancer modifications. Surprisingly, however, the data show a lesser, but still highly significant basal level of GRHL2 mediated enhancer activation in the absence of KMT2C/D. These results collectively establish GRHL2 enhancer activation including chromatin modifications and transcriptional activation as strongly but incompletely dependent on KMT2C and KMT2D. As such, we propose that KMT2C/D are amplifiers rather than essential cofactors in pioneer TF driven enhancer activation.

## Results

### A synthetic GRHL2 system for rapid induction of target transcription in naive ESCs

Previous studies on GRHL2 in ESCs utilized a GRHL2 expression cassette (TRE-GRHL2) integrated into an engineered *Col1A1* locus via FLP-recombination^30^. Induction of GRHL2 expression was driven by doxycycline (Dox) treatment via an rtTA, expressed from an integrated cassette in the *Rosa26* locus. This two loci system is difficult to apply to different mutant backgrounds and has a ramp-up induction due to the requirement for induced transcription and translation. Therefore, we instead built a construct that fused GRHL2 to an ERT domain which was then integrated in the genome by PiggyBac. Constitutively expressed GRHL2-ERT can then be rapidly triggered to enter the nucleus upon tamoxifen (Tam) administration (Fig.1a).

**Figure 1.**
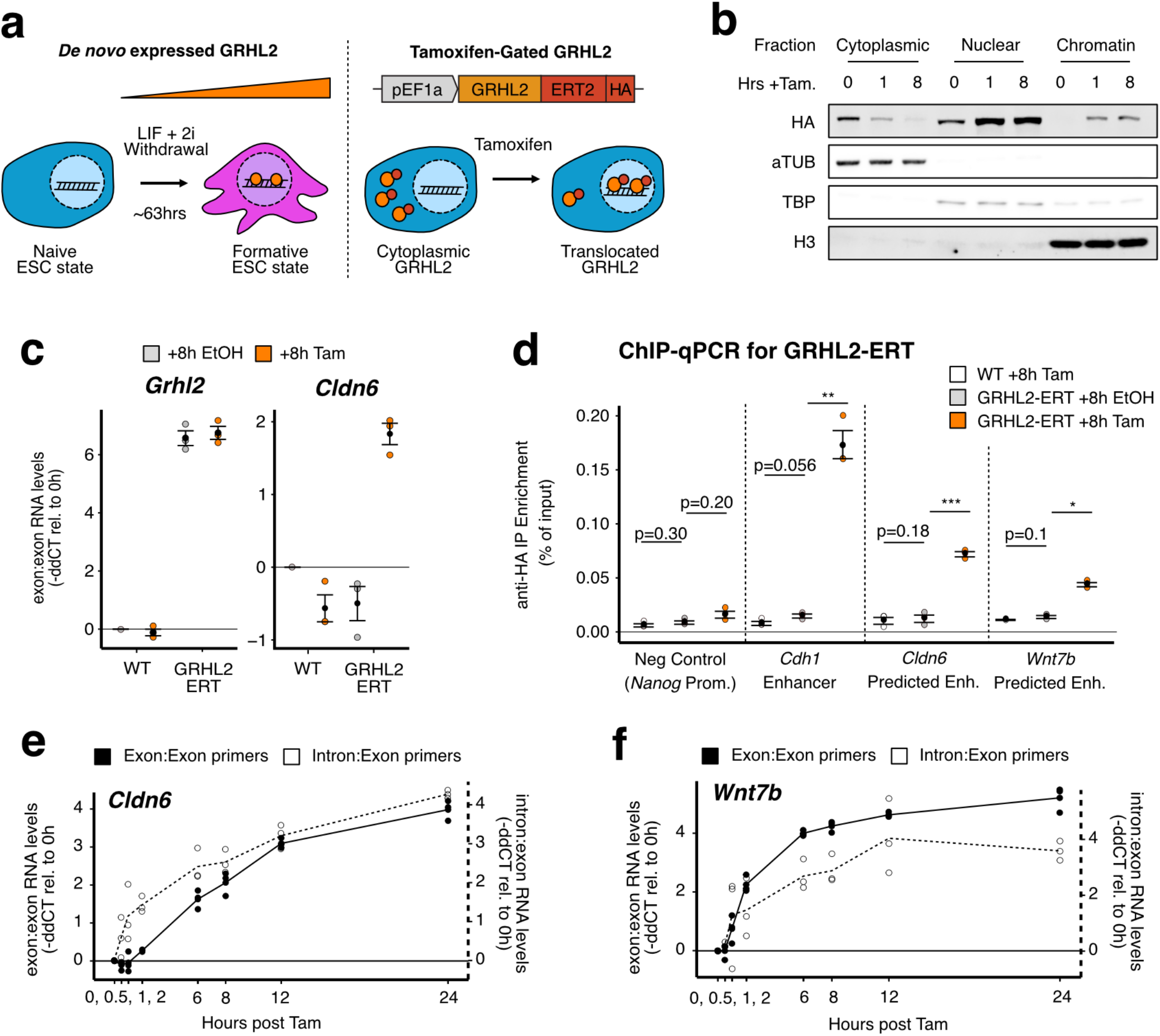
A synthetic system for induction of GRHL2 achieves rapid enhancer binding and gene activation. a) Schematic of the Naive to Formative ESC transition and GRHL2-ERT transgene and conceptual mechanism of induction by tamoxifen (Tam). b) Western blots using subcellular fractions generated from a WT cell line expressing GRHL2-ERT and treated with Tam for 0, 1 or 8hrs. c) qPCR data for Grhl2 and Grhl2 target Cldn6 in WT cells with or without GRHL2-ERT system. Values are -ddCT normalized to GAPDH and relative to WT, 8hr EtOH baseline. Mean and SEM shown in black (n=3). d) ChIP-qPCR for GRHL2-ERT at select enhancers after treatment with Tam. Values normalized by percentage of input. Mean and SEM shown in black (n=3). Paired Student’s T-test with Benjamini-Hochberg correction. e) qPCR measurements over time during Tam treatment. For each, Cldn6 and Wnt7b measurements were made using exon:exon primers (left Y-axis) and intron:exon primers (right Y-axis). Values are -ddCT relative to 0hr induction control. Select statistics shown, all statistics available in Supplemental Table 1. *p<0.05 **p<0.01 ***p<0.001 ****p<0.0001

To first validate the efficacy of the system the engineered ESCs were exposed to Tam for 0, 1, and 8 hours and separated into cytoplasmic, nuclear, and chromatin fractions for evaluation of GRHL2 levels by immunoblotting (Fig.1b). Within 1 hour of Tam treatment GRHL2-ERT is depleted in the cytoplasm and enters the nucleus. The nuclear fraction did show low GRHL2 signal in the absence of Tam but GRHL2-ERT was only detected in the chromatin fraction upon Tam treatment. As an initial test of tamoxifen dependent functional activity of the system, GRHL2-ERT expressing cells and a WT parental line were treated with Tam for 8 hours and collected for qRT-PCR for *Grhl2* and *Cldn6*, a transcriptional target of GRHL2 (Fig.1c). Ethanol (EtOH) was used as a negative vehicle control. Consistent with the enrichment of Grhl2 on chromatin upon induction, Grhl2 was constitutively expressed in the GRHL2-ERT cell line but its target *Cldn6* was only induced upon Tam treatment. No induction was seen in the WT control line.

To further characterize the system, binding was tested at three distinct enhancers. One of the enhancers had been previously validated as a GRHL2 target and is in an intron of the gene *Cdh1*^*31*^. The two others were identified by evaluating the prior dox-inducible GRHL2 data for sites that were bound by GRHL2, gained H3K4me1 and H3K27ac, and induced nearby gene expression upon doxycycline treatment (Fig.S1a,b,c)^31^. One site was near the *Cldn6* gene and the other within an intron of *Wnt7*. Notably, Cdh1 is already highly expressed with broad domains of H3K4me1 and H3K27ac in untreated cells and is not further induced in doxycycline treated cells. In contrast, *Cldn6* and *Wnt7a* mRNA levels were induced 4.6 and 6.7-fold respectively following 24 hours of dox treatment. ChIP-qPCR for GRHL2-ERT at all three sites showed that it bound these sites upon induced nuclear translocation with no evident background in WT or EtOH-treated controls (Fig. 1d).

Given the rapid nature of GRHL2 translocation upon Tam treatment, we next evaluated the transcription kinetics of the system by performing a qRT-PCR time course for *Cldn6* and *Wnt7b*. Measurements were made using both exon:exon primers and intron:exon primers which reflect mature mRNA (spliced) and nascent (unspliced) RNA levels respectively. In both cases, exon:exon and or intron:exon primers, the levels of *Cldn6* and *Wnt7b* were noticeably upregulated within 1 hour after tamoxifen (Fig. 1e,f). Interestingly, while *Cldn6*, intron:exon levels preceded exon:exon upregulation, *Wnt7b* did not, even though *Cldn6* has similarly few introns and thus splicing events. Levels of *Wnt7b* and *Cldn6* continuously increased across 24 hours of Tam and appeared to begin plateauing around 24 hours. Together, these data show that the GRHL2-ERT can rapidly bind chromatin and activate transcription of endogenous target genes in a Tam-dependent fashion.

### KMT2C/D are dispensable for GRHL2 binding but required for downstream enhancer activation

The transactivation domain of human GRHL2 can interact with KMT2C/D^32^. The GRHL2 motif is also highly enriched at formative ESC enhancers that require KMT2C/D for acquiring H3K4me1 and H3K27ac during differentiation from the naive ESC state (Fig. S2a,b,c)^33,34^. Thus, we pursued KMT2C/D as promising candidate coactivators of GRHL2 in activating enhancers upon ESC differentiation. We had previously produced KMT2C/D knockout ESC lines by introducing a cre-ERT into KMT2C-/-;KMT2D fl/fl (conditional knockout, CKO) cells and treating with Tam to produce KMT2C-/-;KMT2D-/- (double knockout, dKO) cells^33,34^. KMT2C is largely absent in ESCs making KMT2D the major contributor to H3K4me and thus we used the parental CKO line as the WT-like control. The GRHL2-ERT system was transposed into the CKO and dKO lines to produce CKO^GRHL2-ERT^ and dKO^GRHL2-ERT^ respectively. Clones showing similar protein levels of GRHL2-ERT were selected for all downstream analyses. (Fig.2a). To avoid the complication of losing KMT2D upon Tam treatment in the CKO cells, all GRHL2 inductions were evaluated at less than 8 hours before KMT2D levels were noticeably impacted (Fig.S2d).

**Figure 2.**
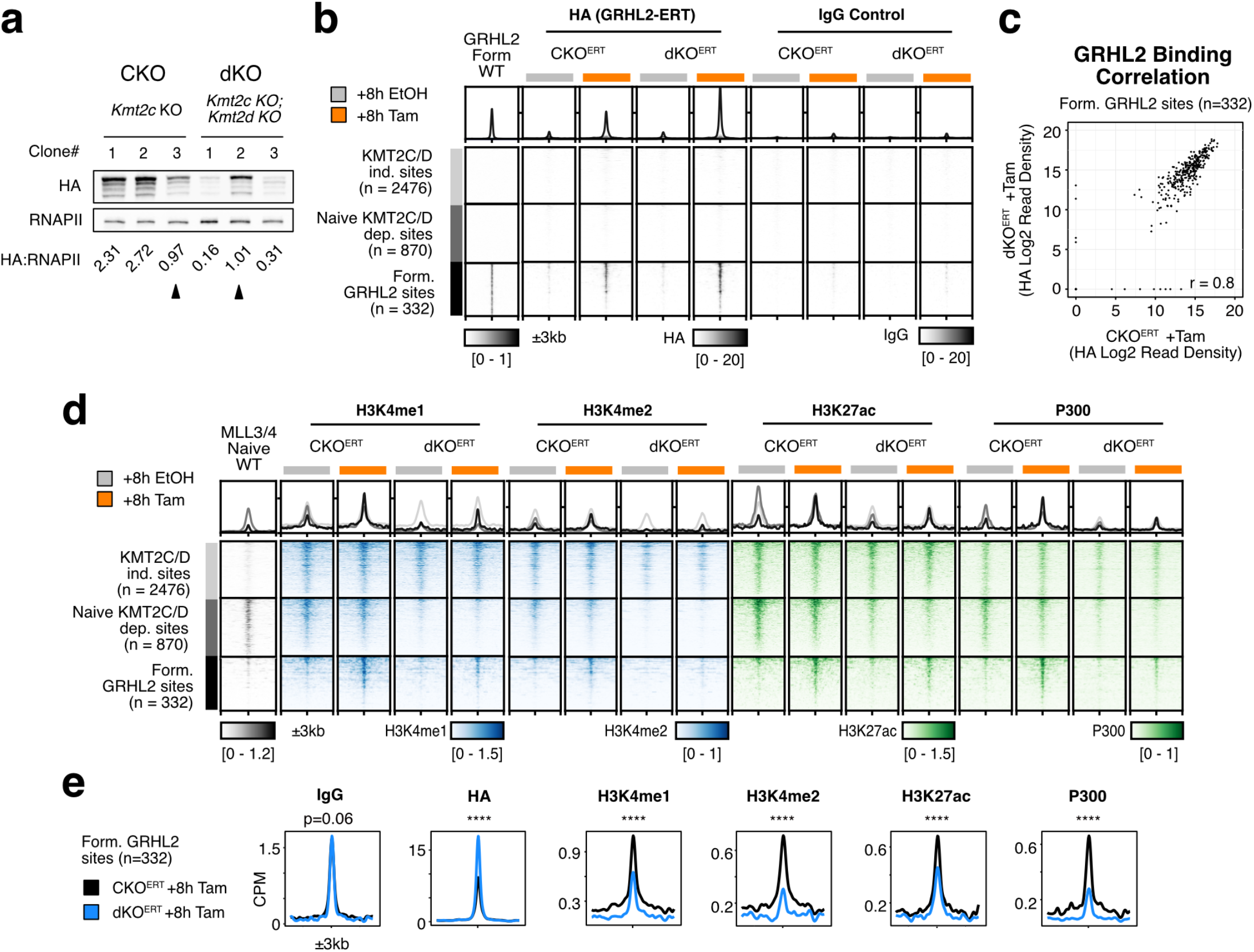
KMT2C/D is required for GRHL2-ERT mediated H3K4me1 and H3K27ac deposition. a) Western blots showing levels of GRHL2-ERT for CKO and dKO clones, downstream experiments conducted using marked clones (triangles). b) Heatmaps of HA and IgG CUT&Tag in CKO^ERT^ and dKO^ERT^ lines with either 8 hours of EtOH (gray) or Tam (orange) treatment. c) Correlation of HA binding signal (Log2 read density) at shared ectopic peaks and Form. GRHL sites for cells treated with Tam for 8h. r = Pearson’s correlation coefficient. d) Same regions as b) but with CUT&Tag for H3K4me1/2, H3K27ac, and P300. All heatmap values are in CPM and the range for each heatmap is the same for each signal profile above. e) Metagene analyses comparing peak signal in CPM of EtOH vs. Tam for CUT&Tag at GRHL2 Form. sites. Wilcoxon Paired Rank Sum test. *p<0.05 **p<0.01 ***p<0.001 ****p<0.0001

GRHL2-ERT binding was profiled by Cleavage Under Targets & Tagmentation (CUT&Tag)^35^ using an antibody toward the HA tag on the GRHL2-ERT fusion protein. IgG was used as a non-targeting control. Heatmaps were generated to visualize and compare GRHL2-ERT binding upon Tam treatment, focusing specifically on 332 GRHL2 binding sites in formative ESCs which had been previously identified by GRHL2 ChIP-seq using GRHL2 knockouts (Form. GRHL2 sites, Fig.2b)^30^. Two additional sets of distal regions were also identified and evaluated as control sets including all distal sites that had either KMT2C/D -independent or dependent H3K4me1/H3K27ac in naive ESCs (see methods for site filtering description). Heatmap analysis revealed a strong enrichment of HA specifically at Form. GRHL2 sites when both CKO^GRHL2-ERT^ and dKO^GRHL2-ERT^ cells were treated with Tam, similar to that observed when plotting published formative GRHL2 ChIP-seq^36^. A slightly greater level of HA signal was found at the GRHL2 sites in dKO^GRHL2-ERT^ relative to CKO^GRHL2-ERT^cells, but the signal was highly correlated (Fig.2c, Pearson = 0.8). These data show that GRHL2 is able to bind endogenous sites independently of KMT2C/D.

To address the functional roles of KMT2C/D at GRHL2 binding sites, we performed CUT&Tag for H3K4me1, H3K4me2, H3K27ac, and P300 (Figs.2d,e). Consistent with filtering, KMT2C/D dependent sites lost H3K4me1/2, H3K27ac, and P300 signal relative to KMT2C/D independent sites. The KMT2C/D dependent sites were also enriched for naive KMT2C/D binding. At Form.GRHL2 sites, few sites appeared to have preexisting signal for each modification/factor and were not enriched for naive KMT2C/D binding suggesting they are not already primed for activation by other factors in naïve ESCs. Upon Tam treatment in CKO^GRHL2-ERT^ ESCs, there were large increases in H3K4me1/2, H3K27ac, and P300 at formative GRHL2 sites. Surprisingly, while this increase was greatly reduced in dKO^GRHL2-ERT^ cells, it was not lost. Similar trends were observed at GRHL2 binding sites in the *Cdh1* and *Cldn6* loci (Fig.S3).

The extent to which GRHL2-ERT bound and impacted sites other than the 332 established formative peaks was also analyzed. To identify additional sites, differential peak analysis was conducted using DiffBind on SEACR^37^ for HA (Figs.S4a,b). In CKO^GRHL2-ERT^ cells, upon Tam treatment 5135 peaks are found to significantly gain HA signal (Log2 FC >1 and FDR <0.05) and 162 peaks with significantly decreased HA signal (Log2 FC <-1 and FDR <0.05). In dKO^GRHL2-ERT^ cells 3013 and 160 peaks showed significant increases or decreases in HA respectively. Upon the intersection of CKO^GRHL2-ERT^ and dKO^GRHL2-ERT^ significantly gained HA peaks and exclusion of Form.GRHL2 sites, 1439 additional “shared peaks” were identified (Fig.S4c). Shared peaks primarily consisted of distal (intergenic or intronic) regions similar to Form. GRHL2 sites (Fig.S4d) and HA signals at these sites were highly correlated between CKO^GRHL2-ERT^ and dKO^GRHL2-ERT^ cells (Fig.S4e). Notably, shared peaks showed lower read density relative to the established Form. peaks suggesting they are lower affinity sites that are revealed upon overexpression of GRHL2. Still, heatmap and metagene analyses showed similar impacts of KMT2C/D loss on H3K4me1, H3K4me2, P300, and H3K27ac at shared peaks as seen with Form. GRHL2 sites (Figs.S4f,g,h). However, unlike Form.GRHL2 sites most shared ectopic sites appeared to have preexisting active enhancer marks prior to Tam treatment.

Together, these data show that within 8 hours of Tam treatment, GRHL2-ERT can access and bind its target sites independent of KMT2C/D. However, KMT2C/D are required for normal levels of H3K4 methylation and H3K27ac deposition.

### KMT2C/D amplifies GRHL2-ERT mediated gene activation

Because KMT2C/D is required to mediate the bulk of active enhancer features at GRHL2 sites, we anticipated that transcription driven by GRHL2-ERT would also be impacted by loss of KMT2C/D. To characterize normal transcriptional activation by GRHL2-ERT and the impact of KMT2C/D loss on GRHL2 targets RNA-sequencing was performed on CKO^GRHL2-ERT^ and dKO^GRHL2-ERT^ cells. Samples were treated with either Tam or EtOH for 8 hours. Differential gene expression analysis was performed using DESeq2^38^ to identify significantly enriched genes in Tam vs. EtOH-treated CKO^GRHL2-ERT^ cells. Upon Tam treatment, 101 genes were upregulated (Log2 FC >0.5 and FDR <0.05) which included *Wnt7b* and other known GRHL2 targets *Cldn6, Cldn4, Epcam, Tmem54, Tacstd2*, and *Acsl5* (Fig.3a). Significant differences in *Cdh1* were not found, as seen similarly in differentiation, and only 15 genes were down regulated. To account for the potential of Tam treatment itself or Cre-ERT activity to impact transcription, Tam treated samples were also collected from the CKO parental line which does not express GRHL2-ERT. Using differential gene analysis to compare the Tam-treated CKO parental line to the EtOH-treated CKO^GRHL2-ERT^ line identified only a handful of significantly impacted genes including 23 upregulated and 6 down regulated genes (Fig.S5a, Log2 FC >0.5 and FDR < 0.05). Importantly, Consistent with immunoblotting (Fig.S2d), a significant decrease in KMT2D transcript was not detected in the CKO parent line within 8 hours of Tam treatment. We found 0 overlapping genes between the 101 upregulated with Tam treatment in the CKO^GRHL2-ERT^ line (Fig.S5a) and the 23 upregulated in the CKO parental line with Tam treatment. Additionally, 71% (72 of 101) genes upregulated by Tam in CKO^GRHL2-ERT^ cells were also significantly upregulated in the previous TRE-GRHL2 lines after 12 hours of dox treatment (Fig.S5b). These data show that the CKO^GRHL2-ERT^ faithfully recapitulates GRHL2-mediated gene induction seen in a WT background.

**Figure 3.**
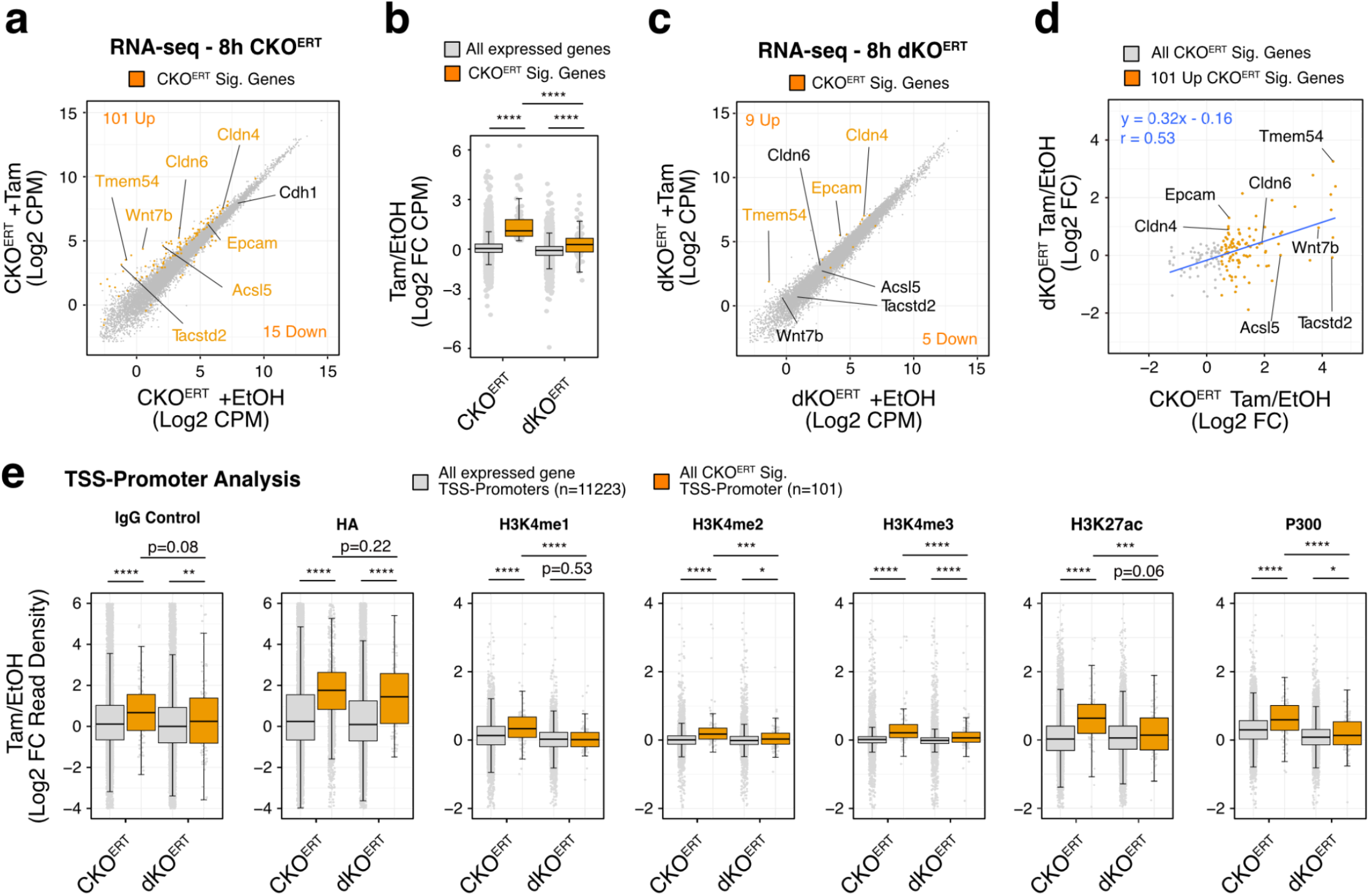
KMT2C/D amplifies GRHL2-ERT mediated gene activation. a) Log2CPM scatterplot of CKO^ERT^ lines treated with 8h of either EtOH or Tam. Significant genes shown in orange (|Log2 FC CPM| > 0.5, Adj. P-value < 0.05). b) Boxplot of Log2 FC CPM (Tam/EtOH) comparing CKO^ERT^ and dKO^ERT^ cells and expression changes at 8h treatment for either all expressed genes (n=12428) or all CKO Sig.genes (n=116). SEM shown. Mann-Whitney U Test, select statistics shown. c) same as Fig.3a, but comparing dKO^ERT^ treated EtOH or Tam for 8h. d) Scatterplot of Log2FC (Tam/EtOH) between CKO^ERT^ and dKO^ERT^ after 8h of treatment. Linear equation represents all CKO Sig. Genes at 8h. e) Analysis of Log2FC in read density for CUT&Tag data at TSSs (+/-1kb) for either all expressed genes (n=11223) or all CKO Sig. genes (n=101). SEM shown. Mann-Whitney U Test, select statistics shown. Complete statistics available in Supplemental Table 1. *p<0.05 **p<0.01 ***p<0.001 ****p<0.0001

Next, we addressed the impact of KMT2C/D loss on GRHL2-ERT mediated transcription by analyzing the fold changes in expression (Tam/EtOH) between CKO^GRHL2-ERT^ and dKO^GRHL2-ERT^ lines for all 101 significantly upregulated genes identified in CKO^GRHL2-ERT^ cells. dKO^GRHL2-ERT^ cells showed a striking decrease, but not complete loss, of transcriptional induction of these genes compared to CKO^GRHL2-ERT^(Fig.3b). Differential gene analysis across all genes in dKO^GRHL2-ERT^ cells (Tam vs EtOH) found 9 significantly upregulated and 5 significantly downregulated genes (Fig.3c). Among those 9 significantly upregulated were GRHL2 targets *Epcam, Cldn4*, and *Tmem54*. Further, scatter plot analysis comparing gene induction following Tam treatment in CKO^GRHL2-ERT^ and dKO^GRHL2-ERT^ cells for the 101 induced genes in CKO^GRHL2-ERT^ showed a correlation between the two, although the degree of induction was dramatically reduced in dKO^GRHL2-ERT^ cells (Fig.3d, Slope = 0.32x, Pearson = 0.53).

We further expanded our analysis to 16 hours Tam treatment. Few additional genes were upregulated in CKO^GRHL2-ERT^ cells at 16 hours compared to 8 hours (135 vs 110) (Fig.S5d). Again, these effects were not secondary to KMT2D loss as only 4 genes were upregulated and *Kmt2d* mRNA levels were unchanged in CKO parental cells at 16 hour Tam treatment (Fig.S5e). Like the 8 hour timepoint, the upregulation of GRHL2 induced genes was significantly reduced, but not lost, in the dKO^GRHL2-ERT^ (Fig.S5f). Differential expression analysis in the dKO^GRHL2-ERT^ identified only 3 genes that were significantly upregulated: *Tmem54, Epcam*, and *Cldn4* (Fig.S5g). Still, correlation analysis of fold induction following 16 hour treatments between CKO^GRHL2-ERT^ and dKO^GRHL2-ERT^ still showed a correlation (Pearson of 0.62), even though the degree of induction was severely blunted (Slope = 0.45x, Fig. S5h). Consistent with these findings UMAP analysis of both 8 hour and 16 hour analysis, showed separation between CKO^GRHL2-ERT^ but not dKO^GRHL2-ERT^ treated with Tam vs. EtOH (Fig. S5i,j).

Changes in transcription are typically highly correlated with changes in several histone modifications at promoters including H3K27ac and H3K4me3^39^. Therefore, to further support results from RNA-seq, we performed CUT&Tag for H3K4me3. These data were combined with the data of other marks described in Figure 2 and evaluated at the promoters [1kb up- and downstream of transcriptional start sites (TSS-promoters)] of the 101 gene upregulated following 8 hours of Tam treatment of CKO^GRHL2-ERT^ cells (Fig.3e). The promoters of all expressed genes (n=12428) were used as a background control. These data showed consistent induction of GRHL2 (HA) binding in CKO^GRHL2-ERT^ and dKO^GRHL2-ERT^ cells. In contrast, induction of all histone marks including H3K4me1, H3K4me2, H3K4me3, H3K27ac, and P300 were reduced. However, aligned with increased transcription, albeit much reduced, all marks were still significantly up or trending up (H3K27ac) in the dKO^GRHL2-ERT^ cells with the exception of H3K4me1. These data show that KMT2C/D are not essential, but are instead amplifiers of GRHL2 driven transcriptional activation.

### KMT2C/D are required for GRHL2 mediated deposition of active enhancer features during formative differentiation

To complement our findings using the GRHL2-ERT system we next assessed the relationship between endogenous GRHL2 and KMT2C/D in the naive to formative ESC transition where GRHL2 naturally becomes expressed *de novo*. We first confirmed using RNA-seq data and immunoblotting on dKO and CKO cells that Grhl2 gained expression during the formative transition (Figs.4a,b). Cleavage Under Targets & Release Under Nuclease (CUT&RUN)^40^ was then used to profile GRHL2 in naive and formative WT and dKO cells. As seen in the GRHL2-ERT system, endogenous GRHL2 binding was not impacted in the dKO cells and showed strong correlation across all 332 sites (Figs.4c,d). We next reevaluated our previously published CUT&RUN data on H3K4me1 across differentiation^41^ and combined it with newly generated CUT&Tag data on H3K27ac and P300 in CKO and dKO cells in the naïve and formative states. Again, consistent with findings with GRHL2-ERT, there was a major reduction in H3K4me1, H3K27ac, and P300 at GRHL2 bound sites in the dKO cells relative to either WT or CKO formative cells (Fig. 4e,f, Fig.S7a). As revealed by metagene analyses, this reduction was highly significant (all factors p<0.0001) despite endogenous GRHL2 binding not being significantly different at these sites (p=0.92)(Fig. S7b,c).

**Figure 4.**
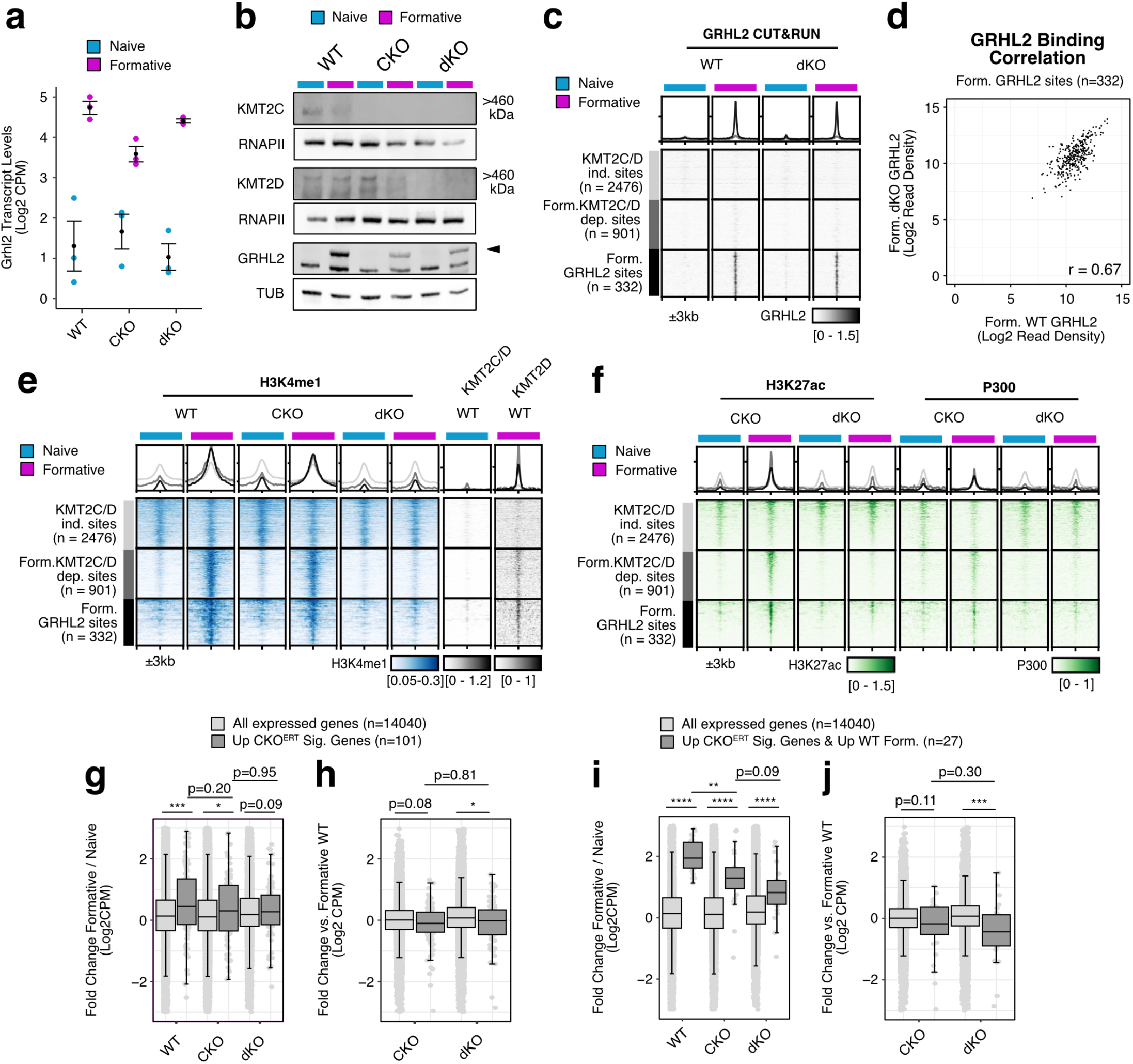
KMT2C/D is required for GRHL2 enhancer activation in the formative state. a) Transcript levels in Log2 CPM of GRHL2 from RNA-seq data. b) Western blots of KMT2C, KMT2D, and GRHL2 in naive and formative ESCs. c) Heatmap visualization of endogenous GRHL2 CUT&RUN data for WT and dKO ESCs in naive and formative states. d) Correlation of GRHL2 CUT&RUN binding signal (Log2 read density) at Form. GRHL sites in Form. WT and dKO cells. r = Pearson’s correlation coefficient. e,f) Same peaks as Fig.4c published or newly generated H3K4me1, H3K27ac, KMT2C/D, and KMT2D data at 332 GRHL2 binding sites (n=332). Data reanalyzed from Boileau et al. 2023 and Dorighi et al. 2018. All heatmap values and range are in CPM. For signal profiles above heatmaps the range in CPM is the same as shown in heatmap for each factor. g,h,i,j) Boxplots of fold change expression between naïve and formative states or formative expression relative to formative WT levels. All boxplots show SEM and all statistical comparisons are Mann-Whitney U Test with Benjamini-Hochberg correction if necessary. Select statistics are shown, complete statistics available in Supplemental Table 1. *p<0.05 **p<0.01 ***p<0.001 ****p<0.0001

Next, we evaluated the impact of the different cell genotypes on upregulation of the 101 GRHL2-ERT targets (from Fig. 3a) during the naïve to formative transition. This set of genes were significantly up during the transition in WT and CKO cells, but not dKO cells (Fig.4g). Consistent with this finding, the fold change in expression of these genes with the transition was significantly reduced in the dKO, but not CKO cells (Fig.4h).

Notably, our previous study revealed that many GRHL2 targets typically undergo expression-neutral enhancer swapping during differentiation. That is, GRHL2 subsumes gene regulation from naïve specific TFs during differentiation as their expression wanes. Naïve TF expression persists into formative dKO cells and could be masking dysregulated GRHL2 targets^41,42^. Thus, we hypothesized we could more precisely assess the role of KMT2C/D on formative GRHL2 targets if we focused our transcriptomic analysis on genes less likely to be regulated by Naïve TFs. Consequently, we filtered our 101 upregulated GRHL2-ERT genes for those genes that are also normally upregulated during the naïve to formative transition, where GRHL2 becomes expressed and functional. Despite filtering resulting in only a small number of genes (n=27), the transcriptional phenotypes were much more striking between dKO and WT cells further supporting the critical role for the KMT2C/D in enabling GRHL2 induction of expression of target genes with the transition (Fig.4i,j). Surprisingly though, there was still significant upregulation of these targets in the dKO background during the transition showing the requirement for KMT2C/D was not absolute. Together, these data are consistent with the GRHL2-ERT system in that KMT2C/D function as critical amplifiers of GRHL2 induction of enhancers and target gene expression during the naïve to formative transition.

## Discussion

In this study we established a synthetic transcription factor system in ESCs by fusing the pioneer factor GRHL2 to an ERT2 domain, enabling rapid and tamoxifen-conditional enhancer binding and activation of GRHL2 target genes. Applying this system to interrogate the functional coactivators of GRHL2 revealed that KMT2C and KMT2D are generally required for H3K4me1, H3K4me2, H3K27ac, P300 recruitment, and transcriptional activation at GRHL2 target loci. However, a basal level of transcriptional activation remained in their absence. Together with complementary analyses on endogenous GRHL2 activity during ESC differentiation, these data identify KMT2C/D as critical coactivators of GRHL2 at enhancers although not absolutely required for its transcription factor activity.

GRHL2 can directly interact with both KMT2C and KMT2D and the two enzymes are required for GRHL2 mediated sensitivity of cancer cell killing by immortalized human NK cells^43^. Here, these relationships are explored more deeply with genomics approaches and in a different cell model, firmly establishing KMT2C/D as general coactivators for nearly all binding sites and targets of GRHL2. This coactivation function is associated with the recruitment of P300 and establishment of H3K27ac at GRHL2 sites. Moreover, transcriptomics data from acute induction of GRHL2 activity shows nearly all changed genes are gaining expression with few downregulated genes. GRHL2 as a transactivator is consistent with work of others^44–46^. Surprisingly though, one group has suggested GRHL2 acts as a repressor inhibiting P300^47^. This difference may represent cellular context or length of stimulation. Our data are based on an acute model and, therefore, almost certainly represent GRHL2’s immediate and direct effects on its targets.

While the CUT&Tag data showed a dramatic decrease in active enhancer modifications in dKO^ERT^ cells, there was still some increase in H3K4me1 and H3K27ac signal at GRHL2 binding sites upon GRHL2 binding. It is challenging to know whether this gain reflects true signal for these modifications or increased background due to increased chromatin accessibility upon GRHL2 binding. If true signal, it would suggest alternative although less robust mechanisms for GRHL2 induced H3K4me1. Likewise, the KMT2C and KMT2D proteins, independent of their enzymatic activity, are generally thought to be critical for P300 recruitment at enhancers^10,34,48,49^. Together with the observation that H3K27ac can be deposited at enhancers without KMT2C/D^41^, our data further strengthens the case for alternative mechanisms for P300 enhancer recruitment. Candidate alternative H3K4 methyltransferases that play a compensatory coactivator role at GRHL2 sites may be KMT2A or KMT2B, where the latter has recently been shown to deposit H3K4me1 at enhancers in redundancy with KMT2C/D^50^. Either way, such findings as well as studies on cooperative relationships between KMT2C/D and other chromatin complexes^48,50–52^ suggest enhancer activation is more collaborative than proposed sequential recruitment models requiring KMT2C/D as essential intermediates.

Consistent with collaborative enhancer activation models, at least some GRHL2 targets were upregulated, albeit less so, in the absence of KMT2C/D. It is unclear what makes these handful of genes different. GRHL2 binding could stabilize other co-bound TFs at specific enhancers which are not KMT2C/D dependent. Candidate TFs which may co-bind and be KMT2C/D independent include GRHL1 and GRHL3 which are both expressed in naïve ESCs and have been shown to heterodimerize with GRHL2^53,54^. The genomic context of GRHL2 binding could also influence KMT2C/D independent activity. That is, chromatin regulator requirements could differ if GRHL2 is bound at or near the promoter versus a distal enhancer in a long-range interaction with a target gene. GRHL2 binding may also facilitate higher order chromatin interactions, bringing multiple enhancers in the vicinity of target gene promoters, independent of the KMT2C/D. Indeed, synergy between multiple distal enhancers has been shown to overcome loss of Cohesin^55^ which is recruited by KMT2C/D^52^. Therefore, such synergy may also overcome the lack of KMT2C/D.

Beyond early development, there are myriad processes where KMT2C or KMT2D loss of function results in impaired cell migration or epithelial regulation and which should now be reevaluated with respect to GRHL2 function (or lack thereof). This includes failures of cells to migrate in gastrulation in *Kmt2c/d* null mice, the human neurocristopathies Kleefstra’s Syndrome (KMT2C-related) and Kabuki Syndrome (KMT2D-related)^56–60^. As well, cancers associated with KMT2C and KMT2D mutations commonly appear in epithelial and ectodermal derived tissues^61^. Whether a KMT2C/D-GRHL2 axis may operate as a tumor suppressing mechanism should be studied in KMT2C/D related cancers.

Finally, based on the depth of mechanistic knowledge available for ESCs and the fast kinetic properties demonstrated for GRHL2-ERT, we envision its application in many more experiments. Rapid induction of GRHL2 will pair especially well with acute perturbation approaches including small molecule inhibitors, PROTACs, or cell lines engineered with degrons. During the course of our own study, another example of a transcription factor fused to an ERT domain was published in cultured mouse pituitary tumor cells^62^. Gouhier et al. leverage the rapid induction of Pax7-ERT to identify the temporal order of enhancer activation for many enhancer features. In this case, a two-step mechanism is found whereby Pax7 primes enhancers first but then requires cell division to dissociate enhancers from lamina and activate them, as indicated by gains in active enhancer marks, KMT2D, Cohesin, and SMARCA4. Most GRHL2-ERT experiments were carried out at 8 hours of induction relative to the 12-hour average cell cycle of an ESC, which would suggest cell cycling is not required by GRHL2 to facilitate enhancer activation. Furthermore, transcriptomics on longer term GRHL2-ERT induction in CKO^GRHL2-ERT^ cells did not show many additional genes being activated, which also contrast with observations with Pax7. Applications of GRHL2-ERT to further study and clarify the mechanistic relationships between chromatin and transcription factors at a sub cell-cycle resolution will be particularly intriguing.

## Materials and Methods

### ESC culture and line generation

Mouse ESCs were cultured in Knockout DMEM (Thermo Fisher, CAT#10829018) supplemented with 15% FBS, L-Glutamine, Penicillin/Streptomycin, NEAA, LIF (1000U/mL), and 2i (1uM MEK inhibitor PD0325901 and 3uM GSK3 inhibitor CHIR99021). GRHL2-ERT proof of principle tests were conducted in WT V6.5 ESCs. Clones used to conduct RNA-seq are derived from WT R1 ESCs. CKO ESCs (KMT2C-/-; KMT2Dfl/fl) and KMT2C/D dKO (KMT2C-/-; KMT2D-/-) are derived from a mixed background of C57BL/6J and 129 strains. The plasmid expressing GRHL2-ERT from the hPGK promoter was integrated into CKO and dKO genomes using cotransfection with a transient plasmid expressing PiggyBac transposase. Transfected cells were treated with blasticidin to select for GRHL2-ERT integration and plated at single cell density in 10cm^2^ plates to isolate and characterize colonies. Formative cells were generated by removal of LIF and 2i. Briefly, 5e4 ESCs were plated per well of a 6 well plate on day −1 in LIF+2i media. To initiate differentiation, LIF and 2i were removed 24 hours after seeding (Day 0). Formative cells were collected on day 3 of differentiation, 63 hours after removal of LIF and 2i. To overcome proliferation defects 7.5e4 KMT2C/D dKO cells were plated per well of a 6 well. Naive cells were passaged and staged appropriately for simultaneous harvest. Lines consistently tested negative for mycoplasm.

### Protein extraction and Westerns

To harvest protein for western assays, cells were trypsinized, washed once with ice cold PBS before adding RIPA buffer with protease inhibitors (Sigma CAT#P8340). After 15 minutes on ice, lysed cells were centrifuged at 16000g for 10 minutes and the supernatants representing whole cell fractions were collected and snap-frozen in liquid nitrogen. Protein quantification was conducted using a Micro BCA protein assay kit (Thermo CAT#23235). 40ug of protein was loaded per well of SDS PAGE gels for western. For subcellular fraction we used the Pierce Subcellular Fractionation Kit (Thermo Fisher, CAT#78840) according to the manufacturer’s protocol. For SDS PAGE of cellular fractions we loaded 20ug per well for each cytoplasmic and nuclear fraction and 10ug for the chromatin fraction.

Westerns were typically conducted using a Bio-Rad system with Tris-Glycine gels purchased from Bio-Rad and transferred to methanol activated PVDF membranes. High molecular weight westerns for KMT2C and KMT2D were conducted using a NuPage SDS-PAGE and Transfer system with an XCell electrophoresis unit according to the manufacturer’s protocols with several modifications (Thermo Fisher, CAT#E10002). In brief, 40μg of whole cell protein lysate was incubated with LDS loading buffer (Thermo Fisher, CAT#NP0007, added BME to 1% final concentration) and incubated at 70°C for 10min. Samples were loaded into 3−8% Tris-acetate gels (Thermo Fisher, CAT#EA03752BOX) and ran in sample running buffer (Thermo Fisher, CAT#LA0041) supplemented with NuPage Antioxidant (Thermo Fisher, CAT#NP0005). Samples were run at 80V for 30min and followed by 120V for 120min. Samples were then transferred to PVDF membranes using NuPage Transfer Buffer (Thermo Fisher, CAT#NP0006, with added 10% Methanol, NuPage Antioxidant, 0.01% SDS) at 30V in a cold room for either 2 h or overnight before blocking.

All membranes were blocked, and stained with primaries and secondaries using Li-Cor Odyssey Blocking Buffer (Li-Cor, CAT#927-60001) mixed 1:1 with TBS. Primary antibodies for western were Rabbit anti-H3 (Cell Signaling, CAT#4493), Mouse anti-alpha Tubulin (Sigma Aldrich, CAT#T-6074), Mouse anti-TBP (Thermo Fisher, CAT#MA1-21516), Rabbit anti-HA (Abcam, CAT#ab9110), Rabbit anti-GRHL2 1:100 (Sigma, CAT#HPA004820), Rabbit anti-KMT2C (provided by Kai Ge), and Rabbit anti-KMT2D (provided by Kai Ge)

### qPCR and analysis

To perform qPCR we first extracted RNA by adding Trizol directly to plates. After adding chloroform, an isopropanol precipitation with GlycoBlue was performed followed by ethanol washes. RNA pellets were resuspended in RNAse-free water and quantified using a NanoDrop. 200ug of RNA was used for cDNA synthesis using Maxima First Strand Synthesis Kit (Thermo Fisher, CAT#K1672) with half reactions according to the manufacturer’s protocol. qPCR on cDNA was performed using SYBRgreen master mix (Applied Biosystems, CAT#A25742) using 6ul final volume on a QuantStudio5 qPCR machine (Applied Biosystems). qPCR primers for targets are listed in Additional File 3. Target Ct values were normalized to GAPDH internally for each sample and then set relative to the naive WT negative control.

### ChIP-qPCR and analysis

Aliquots of 5e6 cells were fixed on a nutator with 1% Paraformaldehyde for 5 min before quenching with 130mM glycine. Cells were washed twice with ice cold PBS then resuspended in RIPA lysis buffer containing protease inhibitors and incubated on a rotator at 4C for 15 min. After centrifugation for 3 min using 2300g at 4C, the supernatant was removed and the pellet was resuspended with 1mL Sonication Buffer (50mM Tris HCl pH8, 10mM EDTA, 0.1% SDS) and incubated on ice for an hour. Sonication was then performed using 1mL milliTUBEs (Covaris, CAT#520135) with a chilled S220 Covaris Sonicator (10% Duty Cycle, Intensity 4, Cycles/burst 200, 6 min total). Successful sonication was performed by visualizing the fragment size of a subset of each sonication using agarose gels a. For ChIP, we quantified DNA concentration using Qubit and used 5μg sonicated DNA diluted to 1mL as input for our immunoprecipitation. 50μL for each target was set aside and stored at 4C as the 5% input control. 950μL of diluted chromatin were mixed with 10ul of Dynabead Protein G beads (Invitrogen, CAT#10003D) pre-bound with 1.8ul of anti-HA (Abcam, CAT#ab9110) and rotated at 4C overnight. IP samples were washed twice with High Salt RIPA (10mM Tris HCl pH8, 1mM EDTA, 0.1% NaDeoxy, 0.1% SDS, 1% Triton X-100, 500mM NaCl), twice with LiCl Wash Buffer (10mM Tris HCl pH8, 1mM EDTA, 250mM LiCl, 1% NP-40, 1% NaDeoxy), and twice with TE Wash Buffer (10mM Tris HCl pH8, 1mM EDTA), with 10 min rotating at 4C for every wash. Beads were resuspended in 50μL SDS Elution Buffer (50mM Tris HCl pH8, 10mM EDTA, 1% SDS) and incubated at 65C overnight shaking at 850RPM along with input control samples. The next day, 0.7μL 10 mg/mL RNAse A and 0.7μL 20 mg/mL Proteinase K were added consecutively and incubated at 37C for 1 hr. Samples were placed on magnet to transfer supernatant and cleaned up using NTB Binding Buffer (Macherey-Nagel, CAT#740595) at a 5:1 NTB:DNA ratio in combination with Nucleospin DNA Cleanup Kit (Macherey-Nagel, CAT#740609) used according to the manufacturer’s provided protocol. DNA was eluted in 50μL EB and used as qPCR input. Resulting Ct values for HA pulldowns and each distal region primer set were then normalized and graphed as a percentage of input controls.

### RNA library generation and sequencing

Total RNA was extracted and purified from cells using Trizol followed by ethanol precipitation. RNA-seq libraries were generated using the QuantSeq 3’ mRNA-Seq Library Prep Kit FWD for Illumina (Lexogen, CAT#A01172) according to their protocol using 200ng of total RNA for input. We utilized the PCR Add-on kit for Illumina (Lexogen, CAT#M02096) to determine an appropriate number of PCR cycles to amplify libraries. Amplified libraries were quantified using Agilent Tapestation 4200. Libraries were pooled and sequenced using a HiSeq 4000 to obtain single end 50bp reads. At least 10 million mapped reads or more per sample were obtained.

### RNA sequencing processing and analysis

Fastq files for RNA-seq samples were processed using Nextflow(Ewels et al. 2020) and the nf-core RNA-seq pipeline v3.9 with default settings. The gene count output from the pipeline was filtered for genes greater than 1 cpm in at least two total samples. Samples were normalized using TMM. Next, Log2 CPM averages were calculated for replicates of each sample for scatterplot visualization and nearest neighbor TSS analysis. All transcriptomics analyses were conducted using CPM values from TMM normalization of all samples except for nearest neighbor analysis where WT and dKO were TMM normalized together. To conduct differential gene expression we performed DESeq2 v1.34.0 analysis using the raw gene count matrix as input for each desired comparison. Gene ontology was performed using ClusterProfiler 4.2.2. Custom R code for other downstream transcriptomics analyses and visualization provided on Github.

### CUT&RUN sequencing and processing

CUT&RUN was conducted using the protocol from Skene et al. 2018(Skene, Henikoff, and Henikoff 2018) with the following modifications: Freshly trypsinized cells were bound to activated Concanavalin A beads (Bang Laboratories, #BP531) at a ratio of 2e5 cells/10ul beads in CR Wash buffer (20mM HEPES, 150mM NaCl, 0.5mM Spermidine with protease inhibitors added) at room temp. An input of 2e5 cells were used per target. Bead-bound cells were then incubated rotating overnight at 4C in CR Antibody buffer (CR Wash with 0.05% Digitonin, 2mM EDTA) containing primary antibody. We used the following antibodies for CUT&RUN: Rabbit anti-GRHL2 1:100 (Sigma, CAT#HPA004820) and 1:100 Rabbit IgG isotype control (Abcam, ab171870). After primary, we washed 3 times 5 minutes each with cold CR Dig-wash buffer (CR Wash with 0.05% Digitonin) and incubated with pA-MNase (1:100 of 143ug/mL stock provided by Steve Henikoff) for 1 hour rotating at 4C. After MNase binding, we washed 3 times 5 each with cold CR Dig-wash buffer, and chilled cells down to 0C using a metal tube rack partially submerged in an ice water slurry. MNase digestion was induced by adding CaCl2 at a final concentration of 2mM. After 30 minutes of digestion, the reaction was quenched using Stop Buffer containing 340mM NaCl, 20mM EDTA, 4mM EGTA, 0.05% Digitonin, 100ug/mL RNAse A, 50ug/mL Glycogen, and approximately 2pg/mL Yeast spike-in DNA (provided by Steve Henikoff). The digested fragments for each sample were then extracted using a phenol chloroform extraction. Library preparation on samples was conducted using manufacturer’s protocols for NEBNext Ultra II Dna Library Prep Kit (New England BioLabs, CAT#E7645) and NEB Multiplex Dual Index oligos (New England BioLabs, CAT#E7600, #E7780) with the following modifications. We input approximately 10ng of sample for half reactions, we diluted the NEBNext Illumina adaptor 1:25, we used the following PCR cycling conditions: 1 cycle of Initial Denaturation at 98C for 30 seconds, 12+ cycles of Denaturation at 98C for 10 seconds then Annealing/Extension at 65C for 10 seconds, and 1 final cycle of extension at 65C for 5 minutes. Following library preparation, double size selection was performed using Ampure beads and quality and concentration of libraries were determined by an Agilent 4200 Tapestation with High-Sensitivity D1000 reagents before pooling for sequencing.

Fastq files for CUT&RUN samples were processed using Nextflow and the nf-core CUT&RUN pipeline v3.1. In brief, adapters were trimmed using Trim Galore. Paired-end alignment was performed using Bowtie2 and peaks were called using SEACR with a peak threshold of 0.05 using spike in calibration performed using the E.coli genome K12.

### CUT&TAG sequencing and processing

CUT&TAG was conducted using the protocol from Kaya-Okur et al. 2020 with the following modifications: freshly trypsinized cells were bound to Concanavalin A beads at a ratio of 2e5 cells/7ul beads in CR wash at room temp. We used 2e5 cells as input per sample. Bead-bound cells were then incubated rotating overnight at 4C in CT Antibody buffer (CR Wash with 0.05% Digitonin, 2mM EDTA, 1mg/mL BSA) containing primary antibody. We used the following primary antibodies for CUT&TAG: 1:100 Rabbit anti-HA (Abcam, abxxx), 1:100 1:100 Rabbit anti-H3K4me1 (Abcam, ab8895), 1:100 Rabbit anti-H3K4me2 (Abcam, abxxx), 1:100 Rabbit anti-H3K27ac (Abcam, ab4729), 1:100 Rabbit anti-KAT2B (Abcam, ab), 1:100 Rabbit anti-H3.3 (Active Motif, xxxx), and 1:100 Rabbit IgG isotype control (Abcam, ab171870). After primary, samples were washed 3 times for 5 minutes each using CR Dig-wash buffer and resuspended in 1:100 secondary antibody (Guinea pig anti-rabbit, Antibodies Online #ABIN101961) in CR Dig-wash buffer at 4C for 1 hour rotating at 4C. Samples were then incubated for 1 hour at 4C with 50ul of approximately 25nM homemade pA-Tn5 in CT Dig300 wash buffer (20mM HEPES, 300mM NaCl, 0.01% Digitonin, 0.5mM Spermidine with Roche cOmplete protease inhibitors added). Recombinant Tn5 was purified and loaded with adapters as previously described (Kaya-Okur et al. 2019). After Tn5 incubation, samples were washed 3 times for 5 minutes each with CT Dig300 wash buffer. Tagmentation was then initiated for 1hr at 37C in a thermocycler by adding MgCl2 to 10mM final concentration in 50uL volume. The tagmentation reaction was quenched immediately afterwards by adding 1.6ul of 0.5M EDTA, 1ul of 10mg/mL Proteinase K, and 1ul of 5% SDS. Samples were then incubated at 55C for 2 hours in a thermocycler to denature Tn5 and solubilize tagmented chromatin. After incubation, samples were magnetized and the supernatant was transferred to new wells where SPRI bead purification was performed using homemade beads to select all DNA fragment lengths larger than 100bp. Samples were eluted in 0.1X TE and approximately half of each sample was used for library preparation using NEBNext HIFI Polymerase with custom indices synthesized by IDT. An appropriate number of cycles for each target was chosen to prevent overamplification bias. After amplification, libraries were purified with 1.2x homemade SPRI beads to select for fragments >250bp and eluted in 0.1X TE. Quality and concentration of libraries were determined by an Agilent 4200 Tapestation with D1000 reagents before pooling for sequencing. CUT&TAG samples were processed similarly to CUT&RUN samples using the same Nextflow pipeline.

### Peak Filtering

The Form. GRHL2 sites were obtained from Chen et al. 2019^21^. In brief, the sites were originally identified by performing ChIP-seq in WT and GRHL2KO Formative ESCs. Peaks were called by MACS2 using GRHL2KO as a background peak set to subtract non-specific peaks (--nomodel --extsize 200 -s 50 --bw 200 -f BAM -g mm -B -q 0.05).

Additional peak sets used in this study were derived as in Boileau et al. 2023^12^. First, significantly different SEACR peaks for H3K4me1 and H3K27ac were determined during the naive to formative transition in WT cells. Naive and Formative H3K4me1 in WT cells was contrasted using DiffBind for SEACR peaks. Filter cutoffs were used to identify naive specific (Log2FC <1, FDR < 0.05), formative specific (Log2FC >1, FDR < 0.05), and unchanging H3K4me1 peaks (FDR > 0.1, |Log2FC CPM| <0.7). Similar was performed for H3K27ac peaks but they were additionally subset by those peaks that overlap with ATAC-seq peaks during the transition to reduce non–specific signals. All derived peak sets were recentered using the coordinates of their overlapping ATAC-seq peaks.

To identify shared KMT2C/D independent H3K4me1/H3K27ac peaks we filtered for all non-significantly changing H3K27ac peaks that also overlapped a non-significantly changing H3K4me1 peak. Then we filtered for peaks that did not lose signal for either mark in the absence of KMT2C/D (FDR > 0.1, |Log2FC CPM| <0.7 for both H3K4me1 and H3K27ac).

To identify naive KMT2C/D dependent H3K4me1/H3K27ac peaks we filtered for naive WT ESC specific H3K27ac peaks that also overlapped with a naive WT ESC specific H3K4me1 peak. We then filtered for peaks that lost signal for both marks in the absence of KMT2C/D (naive dKO / naive WT H3K4me1 Log2FC <-1 and naive dKO / naive WT H3K27ac Log2FC <-1).

To identify formative KMT2C/D dependent H3K4me1/H3K27ac peaks we filtered for formative WT ESC specific H3K27ac peaks that also overlapped with a formative WT ESC specific H3K4me1 peak. We then filtered for peaks that lost signal for both marks in the absence of KMT2C/D (formative dKO / formative WT H3K4me1 Log2FC <-1 and formative dKO / formative WT H3K27ac Log2FC <-1).

### Metagene Analysis

To perform metagene analysis we took the output from computeMatrix (deepTools) with 40bp bins over a 3kb region on either side of the center of each region in our peak bed files. Statistical comparisons were made between samples for the mean CPM signal of the 2 center bins of each region (representing the center 80bp of every peak). We then performed a 5 bin moving window average to each region for visualization.

## Supporting information

Supplemental Table 1 - Statistics

## Data and Code Availability

The raw data generated and related processed files in the current study will be available in the GEO Database upon publication or by request beforehand. Published ChIP-seq, ATAC-seq, RNA-seq, microarray, CUT&RUN, and CUT&TAG sequencing data used in this manuscript are derived from GSE98063, GSE212950, GSE117896, and GSE93147.

Custom code generated and used for analysis in our study upon publication will both be archived on Zenodo and available on Github and under the MIT license for open source.

## Acknowledgements

We thank Kai Ge for providing the original KMT2C KO;KMT2D floxed ESC line and antibodies for KMT2C and KMT2D. We would like to thank the Nextflow community for helpful advice on processing of sequencing samples. We would also like to thank Bryan Marsh, Deniz Gökbuget, Brian Deveale, Yin Shen, Dan Lim, and Licia Selleri for helpful discussions throughout the project.

## Funding

This work was made possible by funding from the National Institute of General Medical Sciences (grant no. R01GM122439). Additionally, R.M.B. was personally supported through the ARCS foundation, the UCSF Discovery Fellowship, and the NIH-T32 Predoctoral Training in Developmental Biology grant (grant no. T32HD007470).

## Contributions

Conceptualization: RMB, RB. Methodology: RMB. Validation: RMB. Formal analysis: RMB. Investigation: RMB, KC. Resources: RB. Data curation: RMB. Writing—original draft: RMB. Writing—review and editing: RMB, RB. Visualization: RMB. Supervision: RMB, RB. Project administration: RB. Funding acquisition: RB. All author(s) read and approved the final manuscript.

## Declarations

The authors have no competing interests.

## Supplemental Figure Legends

**Supplementary Figure 1:**
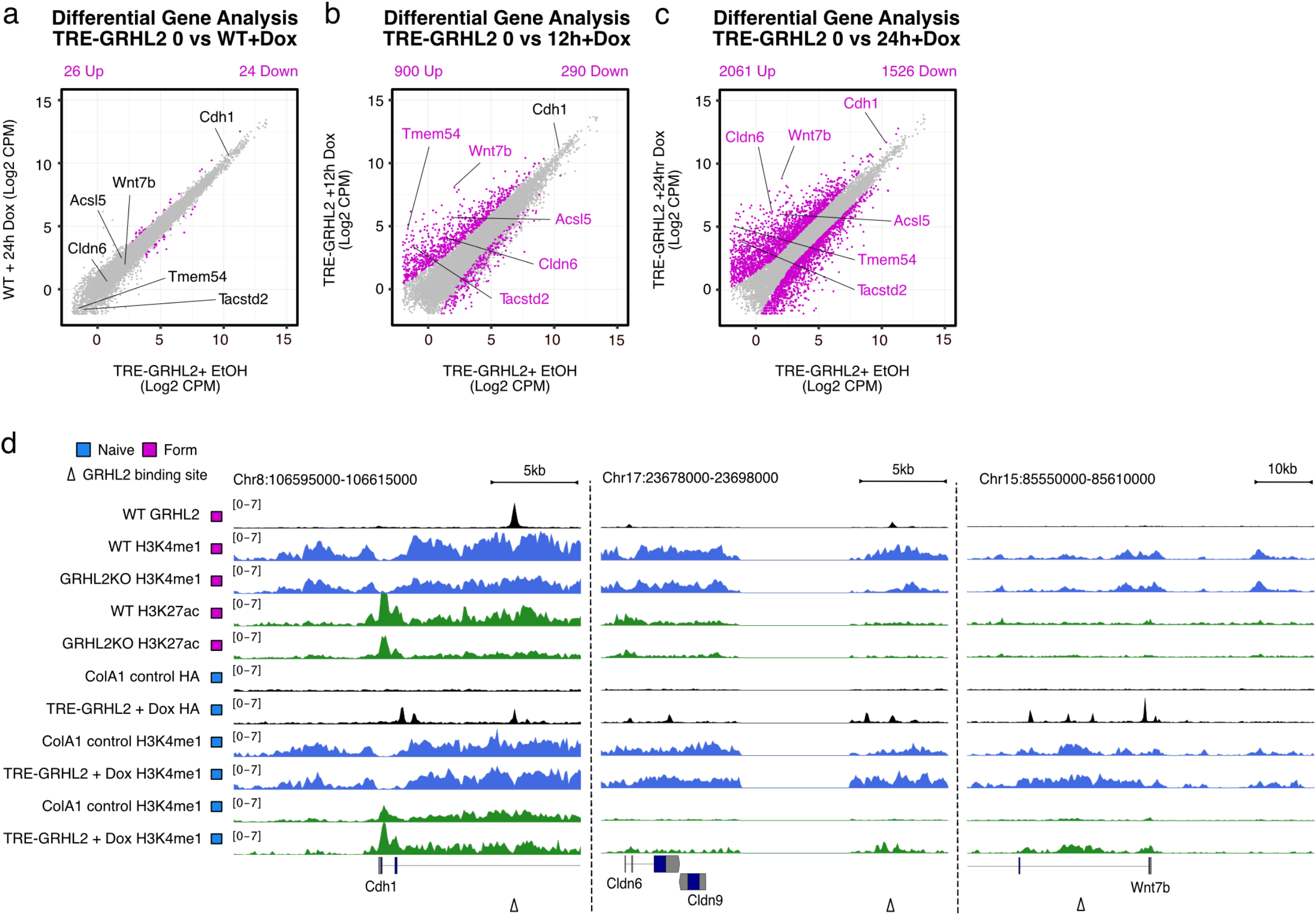
a,b,c) RNA-seq data conducted using the TRE-GRHL2 system at 0,12, or 24 hours dox respectively. Significantly different genes in magenta (|Log2FC| > 1 and FDR<0.05). d) Genome tracks of *Cdh1, Cldn6*, and *Wnt7b* loci. Validated (*Cdh1*) or predicted (*Cldn6, Wnt7b*) enhancers are indicated by arrows. Tracks are scaled by CPM ranges as indicated.

**Supplementary Figure 2:**
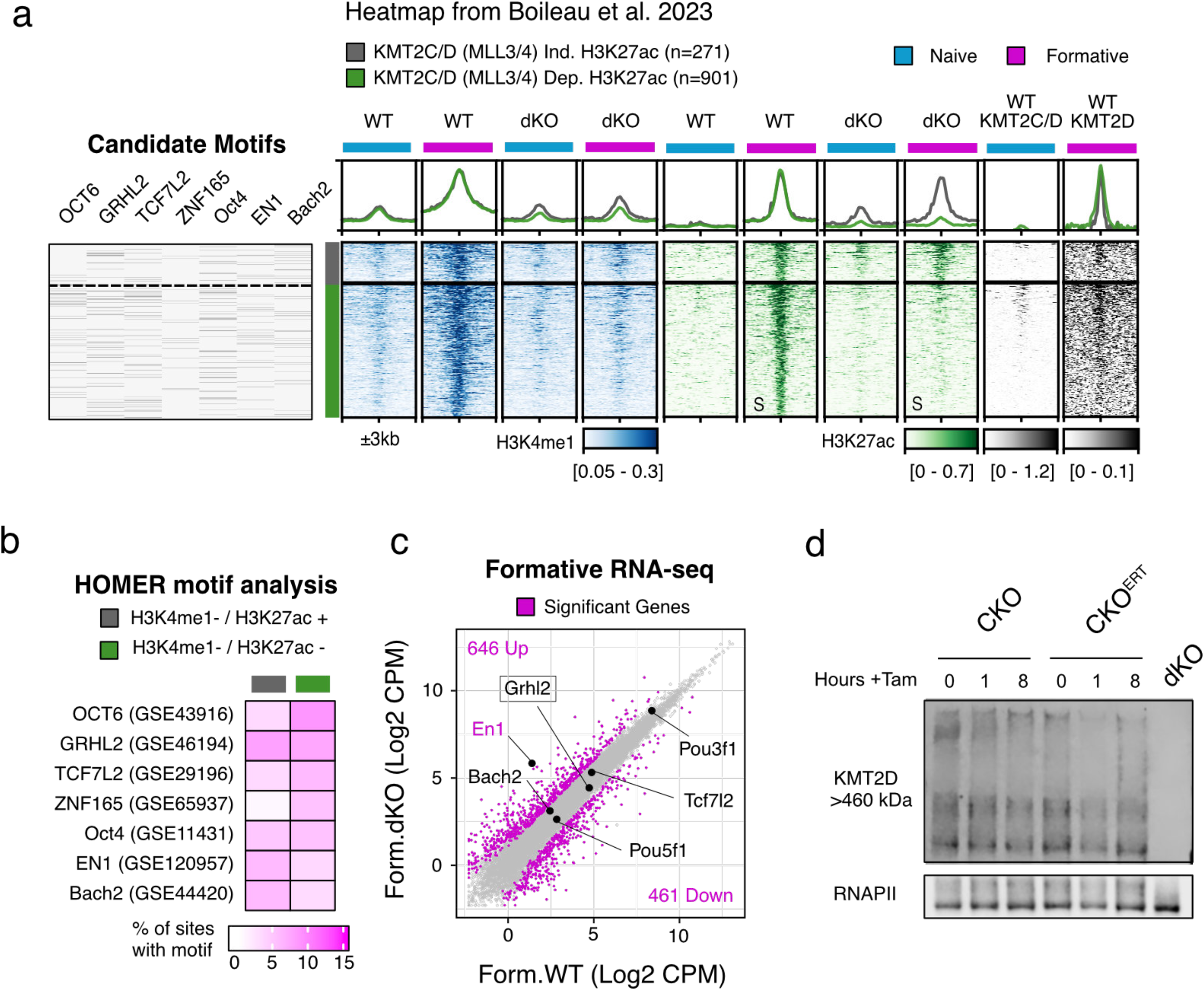
A) HOMER motif analysis conducted on peaks found in Boileau et al. 2023 which are KMT2C/D dependent for H3K4me1 and where H3K27ac is either KMT2C/D independent or dependent. Percent of sites with each motif shown. Positions of motifs are shown sorted similarly to the heatmap. Heatmap values are in CPM and the range used for the heatmaps is the same values used for the metagene analyses above. b,c) RNA-seq data derived from Boileau et al. 2023 comparing Log2 CPM values WT vs. either CKO or dKO cells. Significant genes (|Log2FC| >1 and FDR < 0.05) are in magenta. d) Westerns of KMT2D in either CKO parental or CKO^ERT^ cells treated with Tam for 0, 8, 16 hours.

**Supplementary Figure 3:**
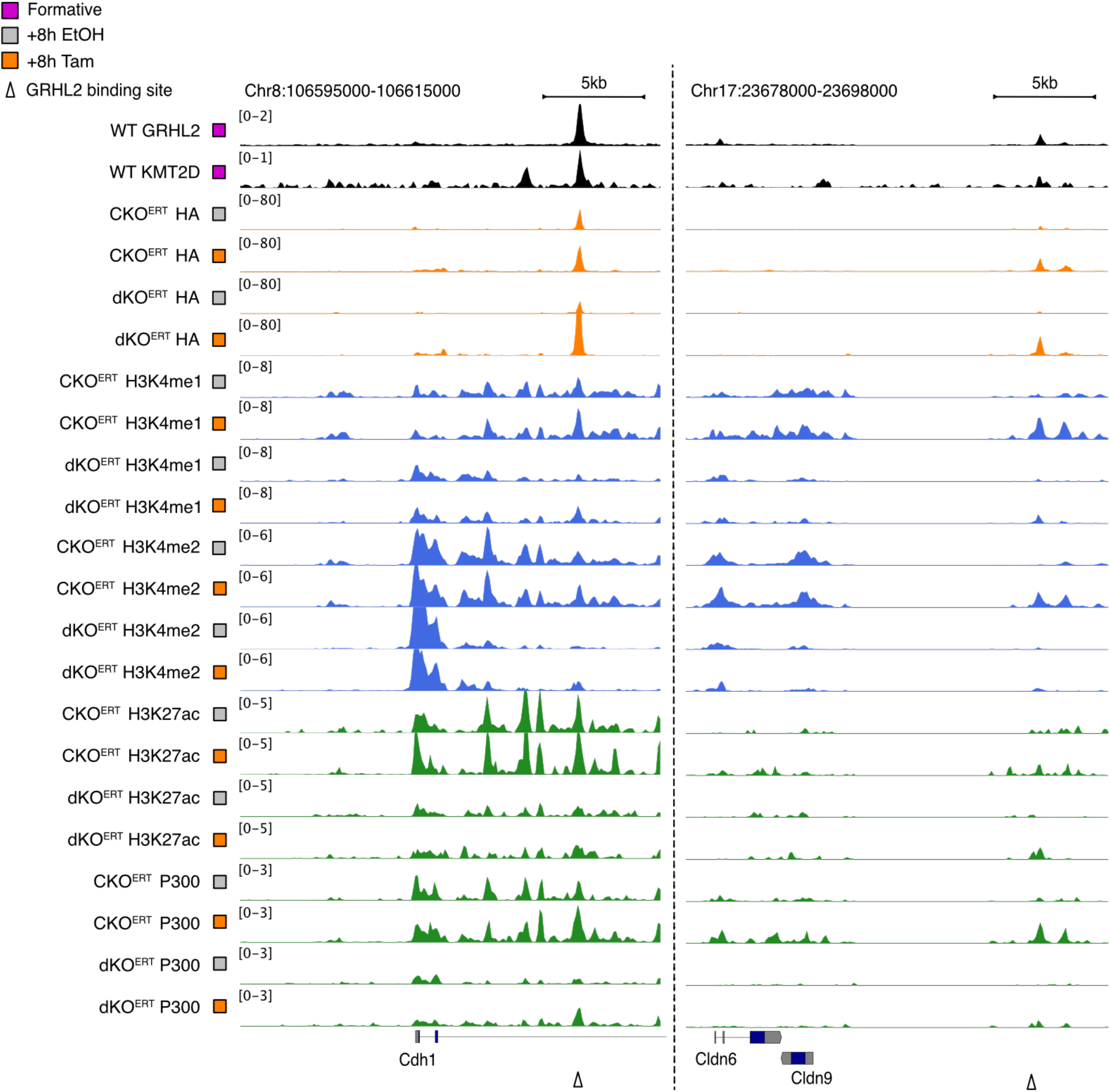
Genome tracks of *Cdh1, Cldn6*. Validated (Cdh1) or predicted (Cldn6) enhancers are indicated by arrows. In dKO^ERT^ cells, *Wnt7b* does not appear to be bound by GRHL2-ERT, unlike CKO^ERT^ cells (not shown). Tracks are scaled by CPM ranges as indicated.

**Supplementary Figure 4:**
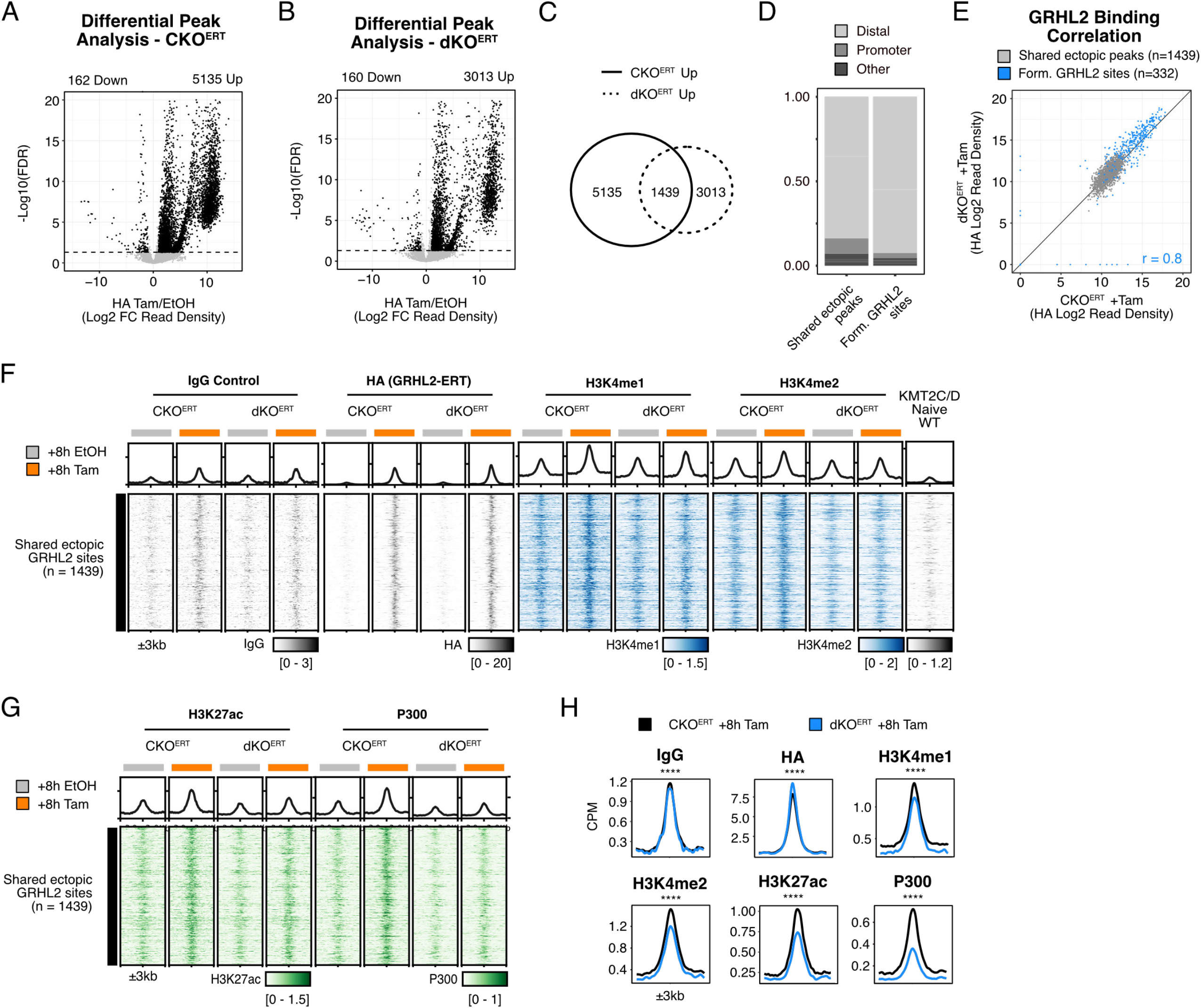
a,b) Differential peak analysis on HA CUT&Tag in either CKO or dKO cells comparing Tam treatment to EtOH controls. Significant peaks in black (|Log2FC > 1| and FDR < 0.05). Dotted line represents FDR = 0.05. c) Comparing peak sets identified in differential analysis to published Form. GRHL sites (top) or with each other to identify shared ectopic peaks excluding Form. GRHL2 sites (bottom). d) Annotation of genomic features for each peak set including distal (intergenic and intronic), promoter, and other (TTS, exon, UTRs). e) Correlation of HA binding signal (Log2 read density) at shared ectopic peaks and Form. GRHL sites for cells treated with Tam for 8h. r = Pearson’s correlation coefficient. f,g) Heatmaps of CUT&Tag data at shared ectopic sites (n=1439) for CKO^ERT^ and dKO^ERT^ lines with either 8h of EtOH (gray) or Tam (orange) treatment. All heatmap values are in CPM and the heatmap range is the same for each metagene analysis above. h) Metagene analysis comparing peak signal EtOH vs. Tam for CUT&Tag at shared ectopic sites. Wilcoxon Paired Rank Sum test. *p<0.05 **p<0.01 ***p<0.001 ****p<0.0001

**Supplementary Figure 5:**
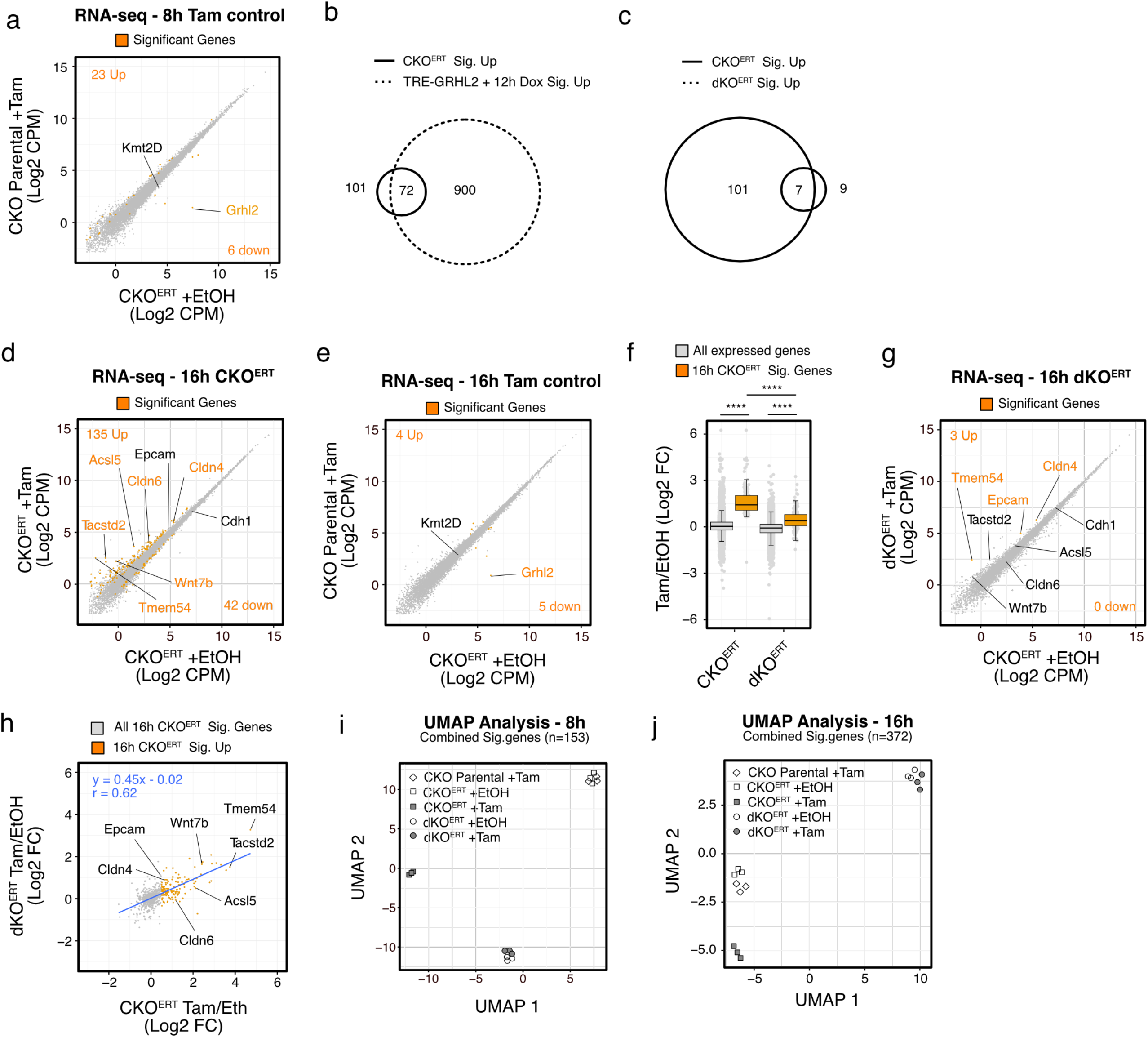
a) Log2CPM scatterplot of Tam control experiment, 8h Tam treatment on CKO cells without the ERT system vs. CKO^ERT^ lines treated with EtOH. b) Overlap between genes significantly upregulated for either CKO^ERT^ Tam/Eth or TRE-GRHL2 (from Fig.S1). c) Overlap between genes significantly upregulated for either CKO^ERT^ Tam/EtOH or dKO^ERT^ Tam/EtOH. d) Log2CPM scatterplot of Tam control experiment, 16h Tam treatment on CKO cells without the ERT system vs. CKO^ERT^ lines treated with EtOH. e) same as Fig.s3a and Fig.3c but comparing CKO^ERT^ Tam/EtOH or dKO^ERT^ Tam/EtOH at 16h treatment. g) Scatterplot of Log2FC (Tam/EtOH) between CKO^ERT^ and dKO^ERT^ after 16h of treatment. Linear equation represents linear modeling using All CKO sig. Genes at 16h. r = pearson’s coefficient. h) Boxplot of Log2FC (Tam/EtOH) comparing CKO^ERT^ and dKO^ERT^ cells and expression changes for either all expressed genes (n=12428) or all CKO Sig.genes (n=177) for 16 hours of treatment. SEM shown. Mann-Whitney U Test, select statistics shown. Complete statistics available in Supplemental Table 1. i,j) UMAP clustering analysis using all 153 significant genes detected in Fig.3a,c and Fig.S5a for 8 hours of treatment or 372 genes total detected in Fig.S5d,e,g for 16 hours treatments. *p<0.05 **p<0.01 ***p<0.001 ****p<0.0001

**Supplementary Figure 6:**
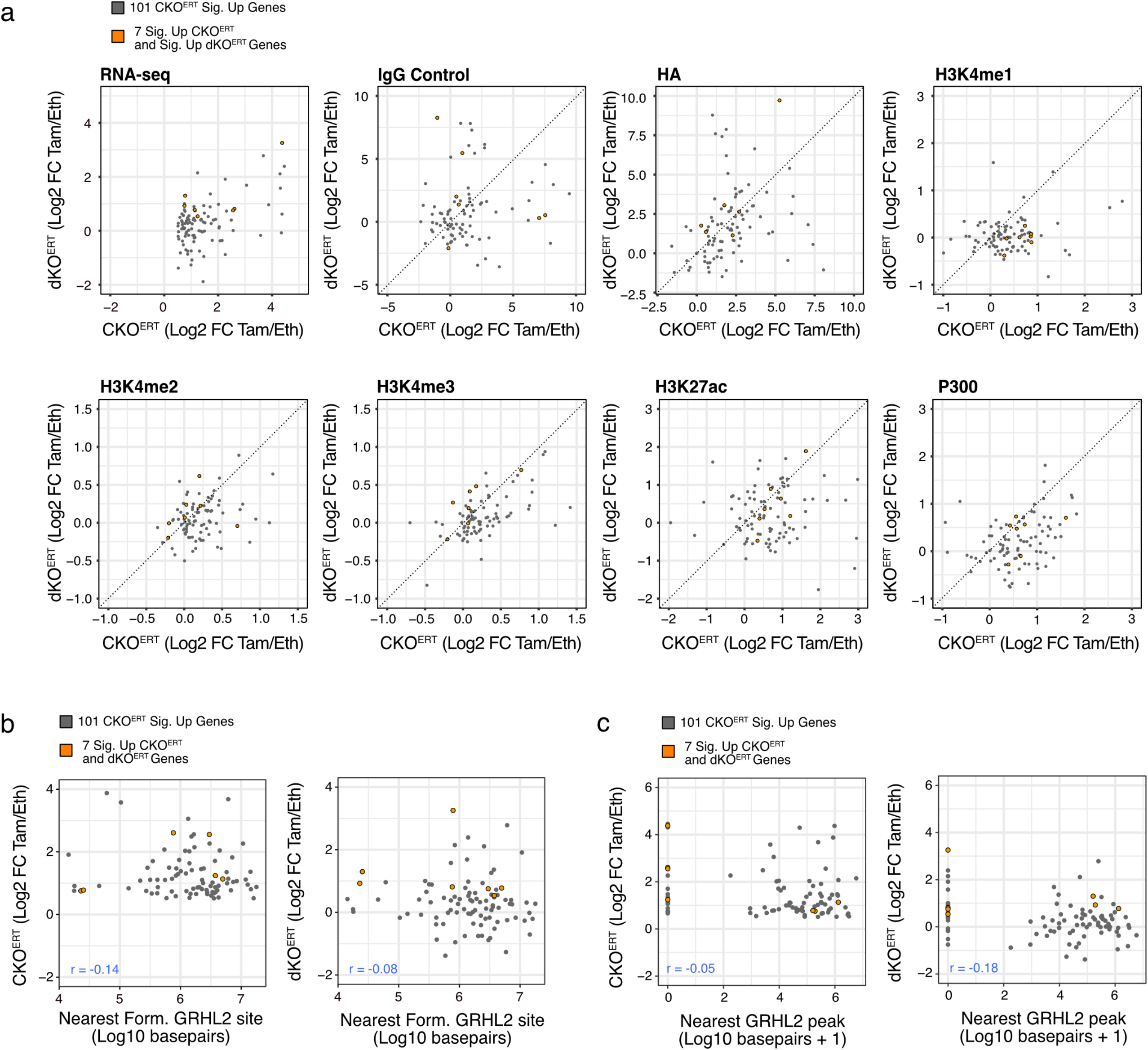
a) Scatterplots showing all significantly upregulated genes in CKO^ERT^ cells (gray) and subset by those that are also significantly upregulated in dKO^ERT^ cells, Log2FC (Tam/EtOH) for each factor. b) Scatterplot comparing fold change gene induction (Log2FC Tam/EtOH) for each cell line to the basepair distance (Log10) to the nearest of 332 Form. GRHL2 sites. c) Same as b, but showing nearest of any GRHL2 peak (shared peaks from Fig.S4). r = Pearson’s correlation coefficient.

**Supplementary Figure 7:**
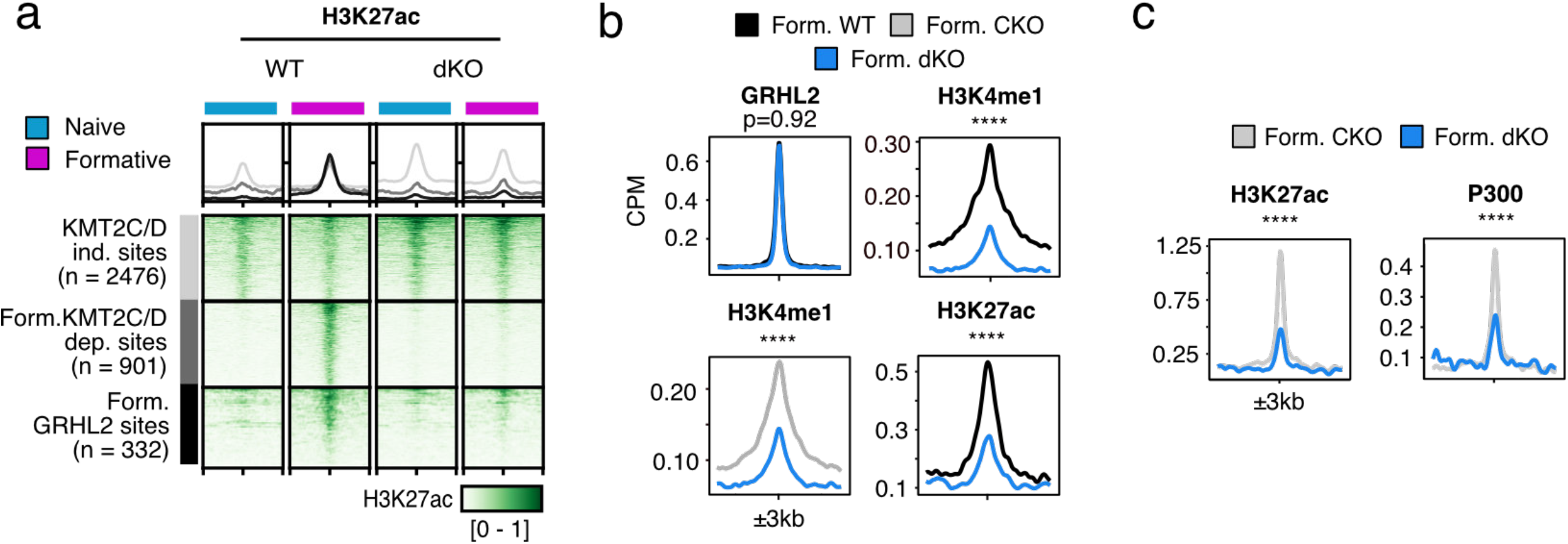
a) Heatmap visualization of published H3K27ac CUT&Tag data for differentiations of WT and DKO cells, Boileau et al. 2023. b,c) Metagene analyses comparing peak signal in CPM of formative WT or CKO vs dKO cells for CUT&RUN and CUT&Tag data at GRHL2 Form. sites. Wilcoxon Paired Rank Sum test. All heatmap values and range are in CPM. For signal profiles above heatmaps the range in CPM is the same as shown in heatmap for each factor. *p<0.05 **p<0.01 ***p<0.001 ****p<0.0001

## Notes

### Competing Interest Statement

The authors have declared no competing interest.

### Summary of Updates

Abstract used in bioRxiv homepage has been revised to reflect abstract in the provided manuscript.

